# A Vision for Ubiquitous Sequencing

**DOI:** 10.1101/019018

**Authors:** Yaniv Erlich

## Abstract

Genomics has recently celebrated reaching the $1000 genome milestone, making affordable DNA sequencing a reality. With this goal successfully completed, the next goal of the sequencing revolution can be sequencing sensors miniaturized sequencing devices that are manufactured for real time applications and deployed in large quantities at low costs. The first part of this manuscript envisions applications that will benefit from moving the sequencers to the samples in a range of domains. In the second part, the manuscript outlines the critical barriers that need to be addressed in order to reach the goal of ubiquitous sequencing sensors.

## Introduction

The cost of DNA sequencing has plunged orders of magnitude in the last 25 years. Back in 1990, sequencing one million nucleotides cost the equivalent of 15 tons of gold (adjusted to 1990 price). At that time, this amount of material was equivalent to the output of all United States goldmines combined over two weeks. Fast-forward to the present, sequencing one million nucleotides is worth about 30 grams of aluminum. This is approximately the amount of material needed to wrap five breakfast sandwiches at a New York City food cart. As a result of the breathtaking pace of advancements, DNA sequencing has become the ultimate back-end of a wide spectrum of biological assays (Shendure and Aiden, 2012). A growing number of these assays simply take advantage of DNA or RNA for labeling, creating molecular post-its that can be read in a highly parallel and cost effective manner (Levy et al., 2015; Zador et al., 2012). Paraphrasing the famous quote by James C. Maxwell (Maxwell, 1864), the aim of many experimental techniques is to reduce the problems of nature to the determination of DNA sequences.

So what is the next frontier of DNA sequencing? For decades, an affordable sequencing platform – the $1000 genome – has been the main focus (Green et al., 2011). While price was under strong selective pressure, the community has largely accepted sequencing devices in any shape and size. Examples of such devices include the now obsolete Heliscope by Helicos Biosciences, which contained a more than 100kg granite slab to stabilize the sequencer (Davies, 2010) and the 860kg Pacific Biosciences’ RSII instrument with its large footprint (**Figure 1A**). In stark contrast, the last year has witnessed the emergence of small footprint sequencers with the successful early access program of the handheld Oxford Nanopore MinION (**Loman and Watson, 2015**) (**Figure 1B**) and an ongoing development of relatively small sequencers by other companies such as Genapsys (**Figure 1C**).

**Figure 1:**
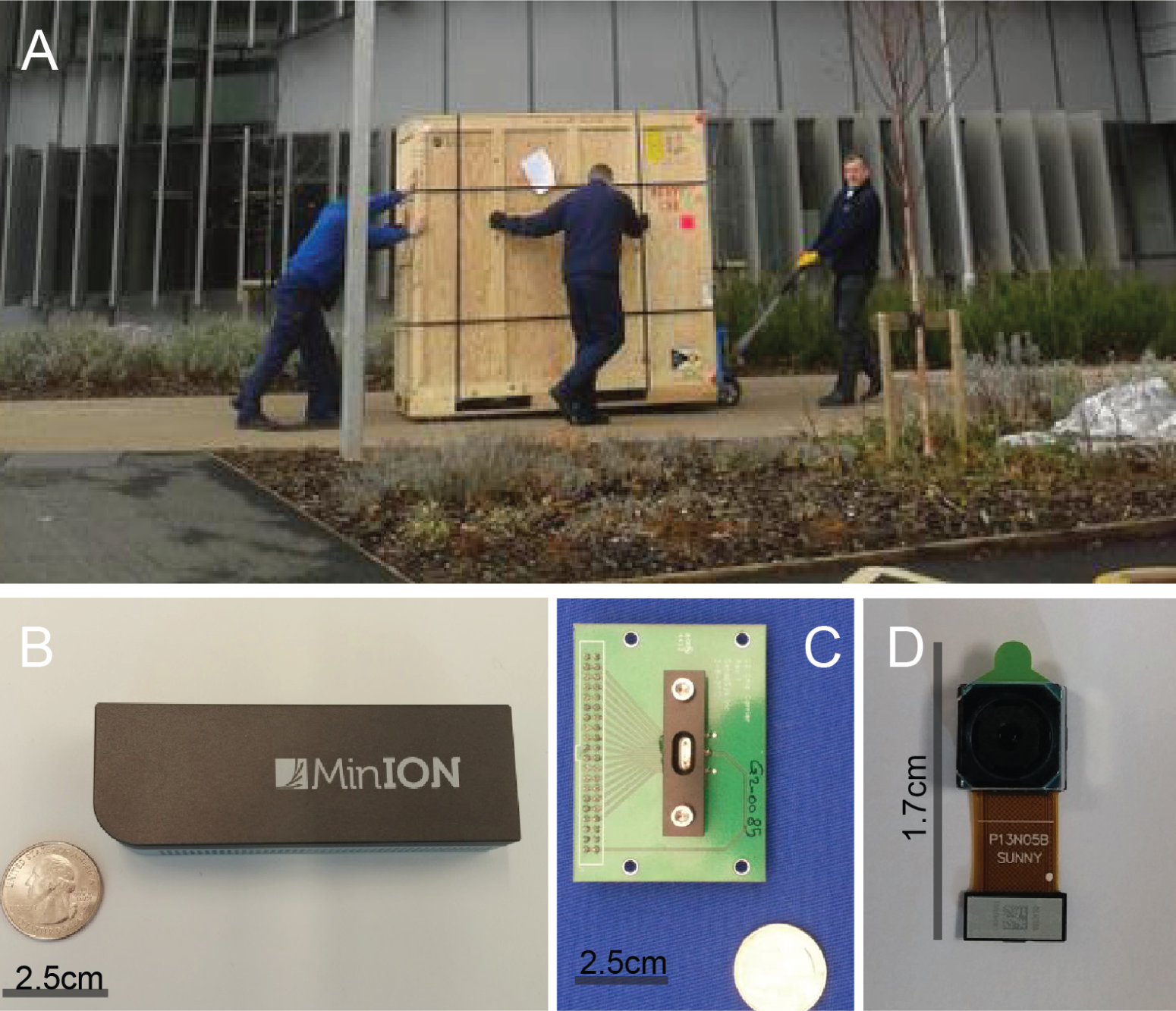
Sizes of sequencing platforms vs. sequencing sensors. For other differences see **Table 1**. (A) Three men haul a 860kg Pacific Biosciences RSII, a sequencing platform, to the University of Exter [photo curtosy of @PsyEpigenetics]. (B) MinION sequencing sensor. (C) An early prototype of a Genapsys flowcell. The company develops an iPad size sequencer. (D) A commodity digital camera chip ready for cellphone integration. Can DNA sequencers be that small?

With these exciting new developments, the next phase of the sequencing revolution is the emergence of DNA sequencing *sensors*. Different from massive sequencing *platforms* such as the Illumina X Ten and Pacific Biosciences RSII, sequencing sensors will be extremely miniaturized devices that include automatic sample preparation with the aim of real-time sequencing in the field (**Table 1**). By integrating these miniature sequencers in larger systems, they will add a DNA-awareness layer to various devices (**Figure 1D**). That being said, sequencing sensors will not replace sequencing platforms in the foreseeable future; the latter will certainly be around for heavy-lifting tasks where scale matters, such as whole genome sequencing. Rather, sequencing sensors will enable a new range of applications that can substantially benefit from moving the sequencers to the samples instead of the traditional way of moving the samples to the sequencers. With this goal in mind, the cost per base pair (bp), extremely high accuracy, and other traditional parameters of sequencing performance will be of secondary importance. More attention will be devoted to the cost of the device, seamless sample preparation, latency of sequencing results, and environmental robustness. Here, I envision potential applications of sequencing sensors in different domains, map chief engineering challenges, and briefly discuss potential regulatory issues of ubiquitous sequencing.

**Table 1:** Sequencing platforms vs. sequencing sensors

| Feature | Sequencing platform | Sequencing sensor |
| --- | --- | --- |
| Operation | Static | Mobile |
| I/O | Fancy touch screens | API |
| Cost of sequencer | Less important | Important |
| Cost per bp | Important | Less important |
| Capacity | Important | Less important |
| Speed | Less important | Important |
| Accuracy | Important | Less important |
| Bandwidth | High | Low |
| Latency | High | Low |
| Main application | Finding genetic variants | Quantifying DNA molecules |
| Made for | Human | Machines |

## Potential applications of sequencing sensors

### Sequencing at home

Sequencing sensors will enable the advent of DNA-aware home appliances, continuing the growing trend of smart devices that monitor the household environment such as Nest Protect that senses CO and smoke levels. Multiple appliances could benefit from integration with sequencing sensors, including air conditioning or the main water supply to monitor harmful pathogens. However, of all possible options, toilets may offer the best integration point. First, toilets regularly collect a large amount of biological material as part of daily life. There is no need for new routines or special procedures, solving a major usability barrier. In addition, it is also possible to monitor pathogens in the water supply. Second, toilet integration can solve certain engineering complications, such as access to fresh water and a place to drain sequencing waste (although access to electricity is expected to be complicated). Most toilets are designed to have space below the water tank, which is rarely occupied. This could be a convenient area to place a small system without substantial miniaturization burden. Third, as opposed to most appliances such as refrigerators or dishwashers, many houses have more than one toilet, increasing the potential market size for such a device. A smart toilet system can convey extremely rich information about household members for medical and dietary purposes. It will allow monitoring of fluctuations in the host microbiome and other potential indicators of the wellbeing of the gastrointestinal and renal systems. Since waste also contains host DNA, it will be possible to seamlessly assign the sequencing results to a specific individual without additional user involvement. In addition, it might be possible to track food composition from waste in order to automatically log dietary habits.

Another exciting opportunity is bringing sequencing sensors to the hands of the general public as a standalone device. With the growing interest in self-tracking, an affordable sequencing sensor with automatic sample preparation has the potential to be a trendy – and admittedly quite geeky – technological accessory. Just a year ago, about 13,000 Kickstarter backers pledged over $2 million to develop SCiO, a low-cost handheld spectrometer that is intended to sense the chemical makeup of food and drugs, highlighting the potential market for similar DNA-aware devices. Beyond the educational advantages, bringing sequencing sensors to the masses can also assist crowd-sourcing DNA or RNA signatures from various sources. This can facilitate the development of interpretation apps in domains that have received relatively little attention from traditional scientific endeavors such as collecting microbiome signatures at various spoiling stages of food or mislabeled food products.

### Health related applications

Real time DNA sequencing has remarkable potential for diagnosing infectious diseases by bringing the sequencers to at-risk individuals. For example, rapid sequencing at airport checkpoints might be useful to control pathogen outbreaks and offer medical assistance to affected passengers. Similarly, a portable sequencer will enable physicians to provide more accurate diagnoses in the field during humanitarian crises or in the clinic without the need to waste time by sending samples to a lab.

A more bold approach is to leverage the at-home sequencing devices presented above. By careful development of at-home tests, sequencing sensors will allow patients to obtain data on infectious disease in the comfort of their own homes, reducing both the misery of getting out of bed while sick and the risk of infecting others. Specialized apps will integrate the sequence reads with other physiological measures and patient complaints. A virtual physician visit (or an FDA-approved fully automated program) will conclude the diagnosis and decide on the course of treatment specific to the pathogen.

Sequencing sensors can also allow pathogen surveillance by constant monitoring of key junctions for disease spreading routes such as hospital laundry (Fijan and Turk, 2012), hotel water systems (Mouchtouri et al., 2007), central air conditioning systems (Tringe et al., 2008), and municipal sewage (King, 2014). Multiple studies have highlighted the utility of DNA sequencing for disease surveillance including the 2011 outbreak of E. coli O104:H4 in Germany (Rohde et al., 2011) and the 2014 Ebola pandemic in West Africa (Gire et al., 2014). These efforts played a key role in identifying the pathogenic strain, tracing its evolution, and reconstructing the spread of disease. However, it should be noted that pathogen surveillance from environmental samples before an outbreak will be far more challenging than these retrospective analyses that had the advantage of diagnosed patients. Recent studies have shown that under certain conditions species annotation pipelines can report questionable results such as the presence of endemic species in samples that were collected far away from their natural habitats or the detection of nearly extinct pathogens (Gonzalez et al., 2014; Afshinnekoo et al., 2015; Mason, 2015). These errors stem from reads with local sequence similarity to the genomes of unrelated organisms or reflect horizontal gene transfer between species. In addition, the field has yet to establish a comprehensive knowledge of the genomic sequences of many microorganisms, increasing the likelihood of species misidentification. Addressing these challenges will necessitate long sequence reads for increased specificity, further maturation of analytical frameworks, and expanding databases of annotated species.

Additional potential usage for DNA sequencing sensors includes integration with medical devices for real-time monitoring of cell-free DNA and RNA in patients. These molecules are released into the plasma due to various biological process, including cancer, conferring a diagnostic vantage point without invasive biopsies (Van Der Vaart and Pretorius, 2008). Multiple lines of evidence have shown that these nucleic acids can be used as biomarkers for a series of acute conditions, such as liver response to drug overdose (Wang et al., 2009), heart attack (Creemers et al., 2012), sepsis (Dwivedi et al., 2012), and brain trauma (Ohayon et al., 2012). Real-time bedside sequencing at relatively short intervals can serve as the basis for a universal apparatus to capture these biomarkers, adding a new layer of information to patient status.

### Forensic and security applications

DNA forensics for counter-terrorism and law enforcement applications can highly benefit from rapid sequencing sensors. Currently, the DNA evidence processing chain includes evidence collection, shipment of evidence to a local crime lab, isolation of DNA, generating an STR profile using capillary electrophoresis, and a database search – a process that usually takes days to complete (Kayser and de Knijff, 2012). Quick sequencing of DNA evidence at the crime scene can reveal the suspect’s identity in the critical window after the event to reduce the chance that the suspect will successfully flee, destroy evidence, or mount an additional attack. First responders of serious crimes will be able to operate the system to automatically notify other forces or border control of the suspect’s identity to better focus efforts to capture the suspect. Such a technology might also have military uses. Recent reports revealed that at least in one instance, US Special Forces used a DNA test for positive identification of a target of high interest (Whitlock and Gellman, 2013). However,despite the importance of the target, the tests took more than eight hours before the assay could provide a final confirmation of the person’s identity.

Another application for rapid DNA-based sequencing is human identification at security checkpoints such as airports. While other biometric signatures such as fingerprints and physiometric properties are accurate and easy to obtain, DNA has the advantage of positive identification in the absence of a person’s inclusion in a database (Erlich and Narayanan, 2014). Technically, this can be done by familial searches that rely on finding autosomal matches with first-degree relatives (Bieber et al., 2006) or by surname inference techniques that leverage Y chromosome matches of distant relatives (Gymrek et al., 2013). With careful implementation that is sensitive to genetic privacy and cultural issues (Kim and Katsanis, 2013), such technology at checkpoints could play a role in fighting human trafficking. This horrific worldwide problem affects millions of victims, most of whom are children or young females and takes a substantial societal and economical toll on certain regions (US Deptartment of State, 2014). Family members of the abducted victim will contribute their own DNA to a specialized database that will only be used to locate relatives, similar to a missing person database. Security forces at airports or borders can check individuals suspected of being at risk. To prevent misuse, samples will be immediately destroyed and results will not be saved upon a negative match.

For a technical perspective, the identification of human DNA requires very low sequencing throughput. Currently, forensic DNA profiles are developed by genotyping 13-20 autosomal Short Tandem Repeat markers, which could be achieved with less than 4000bp of total sequencing throughput (Ge et al., 2012). However, to avoid the lengthy target enrichment, it might be more advantageous to utilize an extremely low throughput shotgun sequencing of the sample. Previous work has shown that any 200 common SNPs in linkage equilibrium create a unique profile for any person on earth (with the exception of monozygotic twins) (Lin et al., 2004). At least theoretically, a similar number of long (> 1000bp) reads could ascertain this number of SNPs. Practically, identification will probably require a few thousand reads to collect additional SNPs in order to obtain higher levels of confidence. The variant calling can be done together with aggressive genetic imputation which was shown to substantially improve the calling of common variants despite extremely low sequencing coverage (Pasaniuc et al., 2012). Finally, sample identification will rely on querying a forensic database of whole genome sequencing or genome-wide arrays.

### Food industry

Rather than having consumers sequence food products at their home, quality control can be integrated at various stages of the entire supply chain. Preliminary studies have shown that DNA sequencing can answer various food-related applications including distinguishing morphologically-similar poisonous and edible mushrooms, detecting the accumulation of pathogenic microbes in meat, tracing the fermentation of cheeses, identifying hidden traces of allergens, and even predicting the ripening time of avocados (Dopico et al., 1993; Galimberti et al., 2013; Wolfe and Dutton, 2015; Dugat-Bony et al., 2015). As such, sequencing sensors can be a highly flexible backend to monitor various issues in the food supply chain. The real challenge, however, will be to perform these assays outside of a laboratory setting.

Instead of analyzing naturally occurring sequences, ubiquitous sequencing in the food industry can also utilize barcoding of food products with artificial DNA tags. Recent work has tested the usage of such tags in the food supply chain by delivering exogenous DNA fragments within a coat of silica, a common food additive, at an extremely low cost. It has been showen that these tags could be used to trace the source of milk in dairy products such as yogurt (Bloch et al., 2014) and serve as a long-term authentication mark of premium olive oil (Puddu et al., 2014). Moving forward, artificial DNA labels can create an edible data structure that includes the type of food, producer, lot number, nutrient information, and presence of known allergens or to simply encode a URL to a webpage that contains this information. The length of the DNA label can be relatively short, as recent DNA-specific encoding schemes have proposed methods to robustly code 1.6bits of information in each DNA base pair (Goldman et al., 2013). DNA-labeled products can establish an inexpensive method to identify ingredients and sources of processed food, tracking authenticity and revealing hidden traces of allergens. In addition, if it is possible to precisely quantify DNA labels from human waste, this scheme coupled with smart toilet systems (discussed above) will enable automatic logging of ingredient consumption. This can revolutionize the tedious manual process of documenting food intake, which is a key component of various diet programs and can help researchers gain substantial amounts of data correlating ingredient consumption and illness. These DNA tags could also label oral medications. Sequencing sensors at toilets could trace the persence of the drug in waste, creaeting a technology to monitor patient adherence regardless of the chemistry of the drug, addressing a major public health issue.

## Technological Challenges

Full realization of DNA sensors will require overcoming a multitude of barriers. Below is an outline of the main obstacles:

### Sample preparation

Mainstream protocols for shotgun sequencing are not compatible with the aim of real time miniaturized DNA sensors. They involve multiple chemical reactions and physical manipulations to purify DNA and tether the molecules to the sequencing apparatus, inducing a substantial latency and complicating miniaturization. Multiplexed target enrichment protocols, such as in-solution capture, are even more involved. They consist of more complex steps, including condensation of intermediate reactions, rapid cooling and heating, and a long (> 24hr) hybridization step.

Recent advancements have the potential to greatly facilitate the advent of rapid and miniature library preparation methods. A new single cell sequencing method has proposed a microfluidic-based technique to perform most of the RNA library preparation within droplets (Macosko et al., 2015; Klein et al., 2015). This procedure compresses a large number of processing stages into one step and creates a scalable method for in-situ library preparation. GnuBIO, a BioRad company, develops a microfluidic-based system for genomic library preparation that is completely integrated within their benchtop sequencer. If successful, this system will allow an end-to-end automated sequencing pipeline that starts with genomic DNA samples. Another alternative is to simplify library preparation by avoiding multiplexed target enrichment. Most of the applications mentioned above could either utilize shotgun sequencing or focus on a single region using a miniaturized PCR machine.

### Supply of reagents

Low maintenance and environmental robustness are key issues for sequencing sensors. Current sequencing technologies require multiple types of reagents such as DNA polymerase (Illumina, IonTorrent, and Pacific Biosciences, Genia), biological pores coupled to a motor enzyme (Oxford Nanopore), modified nucleotides (Illumina & Pacific Biosciences), and exogenous DNA primers (all technologies). These have relatively low durability and some also need to be stored at 4C or −20C. One possible solution to reduce the burden of loading reagents onto sequencing sensors is a user-friendly cartridge on which sensitive reagents can be lyophilized to allow storage at room temperature. A non-mutually exclusive effort is to minimize the types of reagents and prolong their durability. Previous studies have suggested that solid-state nanopores can confer increased robustness and durability over protein-based nanopores in lipid bilayers (Dekker, 2007; Venkatesan and Bashir, 2011). These solid-state nanopores might have the ability to sense naturally occurring DNA without any modification, which can greatly reduce the need for a reagent supply.

### Analytics

An open question is how to perform the data analytics for sequencing sensors. Current sequencing platforms push low level data from the machine and use external computers for almost all of the downstream data processing, from registering the sequencing signals via base-calling to alignment. Mobile sequencing sensors can wirelessly push raw data to the cloud or a local server as they sequence the samples. However, this will require data streams that are sufficiently compact to work with the bandwidth of mobile communication.

Latency is another factor of data analytics. Most sequence analysis algorithms are devised as offline programs that assume that all input data is available at the beginning of the run and that files are read and written at each stage (but see (Chiang et al., 2014)). Latency time could benefit from online bioinformatics pipelines that will align sequence reads as the flow of bases continues to arrive from the basecaller and make on-the-fly decisions before the full data is analyzed, such as pathogen detection from sufficient evidence. Unlike most data analytics today, sequencing sensor analytics should provide succinct and concrete information. The typical user will not care about quality recalibration or left alignment of indels and will have zero knowledge or patience to deal with bioinformatics issues. All he wants are short answers such as “Meat is spoiled: don?t eat” or “suspect identified as Mr. X". Simple and reliable reports will be key.

### ELSI aspects of ubiquitous sequencing

The advent of ubiquitous sequencing can trigger a wide range of ethical, legal, and social implications (ELSI). The general public often considers DNA information as more sensitive compared to other types of personal data (Lunshof et al., 2008; Presidential Commission for the Study of Bioethical Issues, 2012; Rodriguez et al., 2013). This fact is also reflected in certain laws that provide extra privacy protections for DNA information.

To highlight the complexities, consider the recent case of the ‘devious defecator’ (Gilbert, 2015). In this case, a grocery storage company, Atlas, found that one or more of its employees defecated in one of its warehouses, ruining products. The company hired a forensic lab to obtain DNA from the feces and requested a DNA test from two employees thought to be responsible. The employees both sued Atlas since, according to the US Genetic Information Non-discrimination Act (GINA), it is unlawful for employers “to request, require, or purchase genetic information with respect to an employee”. The court ruled in favor of the plaintiffs and awarded $2.25 million in damages although the genetic test was done on CODIS STR markers that do not reveal any health status. Most people are not aware of the challenges associated with analyzing human DNA samples and the democratization of sequencing can expose them to legal liabilities.

Another laws also prohibit the collection of human DNA without consent. For example, in the UK, the Human Tissue Act requires consent for any collection of human DNA if there is intent to perform DNA analysis. The Israeli Genetic Information Law only authorizes genetic labs to perform tests on human DNA, with the exceptions of research and law enforcement purposes. In the US, abandoned DNA is not protected by the federal 4th amendment and it is legally possible to collect and analyze samples from public places (Joh, 2006) as was well demonstrated by a recent art project (Gambino, 2013). However, certain states have imposed further regulation. For example, according to Section 79-L of the New York State Civil Rights Law, it is unlawful to “diagnose the presence” of genetic variations that are linked to human diseases without consent. A California senator recently proposed a more strict law similar to the UK law that will prohibit any collection of human DNA without consent with the exception of law enforcement purposes.

These laws can complicate even benign applications such as environmental tracking of bacterial communities in the sewage system. Previous shotgun sequence-based studies showed that such samples could contain small levels of human contamination. While human identification is probably not easy from these contaminants, their actual analysis could be interpreted as unlawful.

Since it is nearly impossible to avoid collecting human DNA in some settings, it will be beneficial to devise mitigation methods. These can include a warning message to the user that human DNA might be collected as part of the sequencing operation, which might violate local laws. Additional mitigation techniques could rely on a built-in suppression of reads that align to the human genome before analysis or storage. While this option has so far been incomplete (Ames et al., 2015), it will show that a reasonable effort was made to address privacy concerns. These mitigation techniques should be incorporated as part of the sequencer API by setting standard commands such as ‘get consent’ or ‘suppress human DNA’. Such API commands will reduce the burden on application developers and will standardize the mitigation steps. Besides technical mitigation, device manufacturers can facilitate a public discussion about social norms and etiquette on using such sequencing sensors by the general public. Notably, privacy concerns were one of the factors that hampered the adoption of Google Glass, emphasizing the importance of addressing these ELSI issues early.

## Summary and the path foward

With the advent of miniature sequencing devices such as Oxford Nanopore’s MinION, we are on the cusp of truly democratizing DNA information by placing sequencers in the hands of the general public. Ubiquitous sequencing will create an incredibly powerful vantage point for observing massive amounts of DNA and RNA information within its natural context. This will open the possibility of integrating DNA data with other types of sensor information and to obtain a more comprehensive picture of the world around us.

This paper presented a range of potential applications that could benefit from sequencing sensors. But what could be the path forward to map which of these applications would be of interest to the general public? Importantly, most of the applications are in domains that are quite conservative and highly regulated, such as DNA forensics or medical devices. These might be less amenable testing grounds for new technologies. An interesting alternative is to consider citizen developers as an initial focus of sequencing sensors. Such hacker communities have played a pivotal role in the emergence of multiple technologies, from personal computers in the 70’s, Linux in the 90’s, and 3D printing and Rasberry Pi more recently. Similar communities exist in genetic genealogy, where non-scientists have created an impressive set of applications to analyze genetic information (e.g. GedMatch.com). By utilizing the power of citizen developers, it will be possible to test applications and accelerate R&D to mature sensors before testing them in one of the more conservative fields.

In any case, like any powerful technology, sequencing sensors will create a range of societal questions necessitating an ongoing discussion between all stakeholders about the benefits and risks and placing safeguards that will increase public trust in the concept.

## Acknowledgments

I would like to thank Dina Zielinski, Brian Houck-Loomis, Itsik Pe’er, and Shree Nayar for useful discussion. Y.E. holds a Career Award at the Scientific Interface from the Burroughs Wellcome Fund. This study was partially supported by an NIJ award 2014-DN-BX-K089.

### Potential Conflict of Interest

YE serves as on the Scientific Advisory Board of Kailos Genetics and BigDataBio.

